# Identification of a novel *CLPX* variant in a mixed breed dog with anemia and spinocerebellar ataxia

**DOI:** 10.1101/2025.09.15.676223

**Authors:** Bianca S. de Cecco, Jeanna M. Blake, Namju J. Kim, Madeline C. Coffey, Andrea Johnston, Andrew D. Miller, Kari J. Ekenstedt, Jeongha Lee

## Abstract

Spinocerebellar ataxia (SCA) or hereditary ataxia is a progressive neurodegenerative disorder primarily manifesting as cerebellar or spinocerebellar dysfunction, resulting in the loss of motor control and voluntary muscle coordination. SCAs are typically inherited conditions, with causative genetic variants identified in multiple genes in people and across various dog breeds. Recently, an atypical case of SCA was documented in a mixed breed dog. In addition to the classic clinical signs and spinocerebellar lesions of SCA, the dog had retinal and optic nerve degeneration and severe, non-regenerative anemia. Whole-genome sequence (WGS) of the affected dog did not reveal any previously identified canine SCA-associated variants. Subsequent variant filtering against a control cohort of over 700 unaffected dog genomes identified a homozygous 4-base-pair frameshift deletion in *caseinolytic mitochondrial matrix peptidase chaperone subunit X* (*CLPX*) [XM_038580726.1:c.1723_1726del]. *CLPX* encodes a subunit of the ATP-dependent ClpXP protease, a molecular chaperone involved in mitochondrial protein degradation. The variant is predicted to cause a frameshift and a premature stop codon within 17 amino acids, truncating approximately 6.64% of the protein. Our study is the first to explore the association of *CLPX* variants with SCA in any species. Given the high evolutionary conservation of *CLPX*, this report of a *CLPX* variant associated with SCA in a dog may have relevance for understanding *CLPX*-related neurodegeneration and/or anemia in other species.

**Author Summary:** A young mixed-breed dog developed a gait abnormality that progressively worsened, together with vision loss, and severe anemia. Despite treatment, the dog’s condition deteriorated, and he was humanely euthanized. An autopsy revealed extensive abnormalities in the brain, spinal cord, eyes, and bone marrow. These histologic findings supported a diagnosis of spinocerebellar ataxia (SCA), also known as hereditary ataxia, which is a genetic neurological disorder that results in impaired movement and diminished coordination. Genetic analysis identified a previously unreported mutation in the *CLPX* gene. *CLPX* plays a key role in mitochondrial protein quality control by helping break down damaged or misfolded proteins within mitochondria—cell structures critical for energy production that are particularly crucial in high-demand tissues like the brain. This mutation likely disrupted normal CLPX protein function, leading to both nerve damage and impaired blood cell production. While related genes are known to cause similar conditions in humans, this is the first time a naturally occurring *CLPX* variant has been identified in an SCA case in any species. Because *CLPX* is highly conserved between dogs and humans, this finding may offer valuable insights into rare inherited neurological diseases in people.

## Introduction

Spinocerebellar ataxias (SCAs), also called hereditary ataxias (HAs), are a large heterogeneous group of neurodegenerative diseases that have inherited cerebellar or spinocerebellar dysfunction as a core feature (1). SCAs in dogs are characterized by progressive symmetric ataxia, truncal sway, and impaired balance (1). In humans, the classification of SCAs was previously based on neuropathologic features and divided into three categories including olivopontocerebellar atrophy, cerebellar cortical atrophy, and spinocerebellar degeneration; however, in the 1980s with advances in genetic mapping, SCAs were reclassified according to their clinical presentations associated with genetic abnormalities (2, 3). In people, over 40 SCAs have been subtyped, and most of the SCAs are known to be inherited in an autosomal dominant manner due to trinucleotide expansion repeats in genes such as *ATXN1* and *ATXN2* (4). Non-expansion variants in other genes cause SCA in an autosomal recessive manner via toxic RNA gain-of-function, mitochondrial dysfunction, autophagy, transcriptional dysregulation, and/or channelopathies (5). The genetic mapping of these conditions allowed a deep understanding of the disease’s mechanisms and demonstrated that different genetic variants could lead to similar neuropathology (2, 6).

Currently, hereditary ataxias in dogs have been categorized according to their neuropathology. Urkasemsin *et al.* (2014) classified HA into five categories: cerebellar cortical degenerations, spinocerebellar degenerations, canine multiple system degeneration, cerebellar ataxias without significant neurodegeneration, and episodic ataxia. The most recent classification scheme was published by Stee e*t al.* in 2023, which divides HA into four categories: cerebellar cortical degenerations, spinocerebellar degenerations, cerebellar ataxias without substantial neurodegeneration, and multifocal degenerations with predominant (spino)cerebellar component (1, 2). Due to the phenotypic heterogeneity of SCA and ambiguous disease pathogenesis, genetic analysis is crucial in subtyping the disease and understanding the underlying molecular mechanisms. In the last decade, 18 new genetic variants associated with hereditary ataxia have been identified in dogs (1). Many of the variants causing SCA are breed-specific or restricted to a small number of related breeds. For example, a missense *SCN8A* variant is connected to SCA exclusively in the Alpine Dachsbracke breed (7), and an expanded trinucleotide repeat in *ITPR1* is observed in the Italian Spinone (8). Conversely, others are found in multiple breeds, such as the *KCNJ10* missense variant found in Dachshunds, Jack Russell Terriers, Parson Russell Terriers, and Smooth Fox Terriers (9, 10); it is worth noting that additional, different *KCNJ10* variants have been recorded in other breeds with SCA (11–13). The disease pathways identified in dogs share many genes with human cases of SCA, including *ITPR1* (above) and others, such as *GRM1*, *SNX14*, *SPTBN2*, and *CAPN1* (14). Furthermore, findings in spontaneous canine diseases can inform human studies; for example, a truncating *KCNJ10* variant in Malinois dogs demonstrated a new phenotype not previously observed in humans with *KCNJ10* variants (14). Due to these shared genetic mechanisms, dogs are excellent animal models for SCA. The aim of this study is to describe the neuropathology and genetic findings of a case of spinocerebellar ataxia in a mixed-breed dog.

## Results

### Clinical examination

The case dog, a 1-year-and-11-month-old neutered male, reported to be a Collie mix, was presented for progressive weakness and a severe, non-regenerative anemia. The dog had a history of chronic relapsing anemia, which started at 11 months of age. These clinical signs initially responded to prednisone (2 mg/kg/day) but relapsed after treatment. Since his adoption at 6 months of age, he remained the smallest among his littermates and had comparably lower energy, with pelvic limb paresis. The pelvic limb weakness slowly progressed; by 1-year-and-8-months of age, the dog had difficulty standing by himself and was not able to walk on slick surfaces. It was reported that two out of six other littermates had similar clinical signs of hindlimb weakness; however, these other dogs were not available for assessment.

A complete blood count (CBC) confirmed a non-regenerative anemia (Hct 14.2%, reference interval = 35-54%; absolute reticulocyte count 2,500 /µL, reference interval = <92,000 /µL, indicating inadequate or no regeneration; S1 Table) and blood chemistry analysis was unremarkable except for mildly increased AST activity (52 U/L, reference interval = 0-50 U/L). Both the saline agglutination test and 4Dx SNAP test (IDEXX) were negative, decreasing the likelihood of immune-mediated hemolytic anemia and multiple vector-borne infectious diseases (heartworm disease, Lyme disease, anaplasmosis and ehrlichiosis), respectively. Given the previous response to prednisone and the nonregenerative anemia, a presumptive diagnosis of precursor-targeted immune-mediated anemia (PIMA) was made. Given the pelvic limb weakness in both the patient and its littermates, a genetic test for degenerative myelopathy (*SOD-1* variant (15), Animal Molecular Genetics Laboratory, University of Missouri College of Veterinary Medicine) was performed; the result was negative (homozygous wild-type).

On neurological examination, the dog had bilateral absence of menace response, severe pelvic limb ataxia, reduced conscious proprioception in all limbs, and pain on cranial cervical palpation. The dog was weakly ambulatory, and neuroanatomical localization was multifocal, involving the brain and C1-C5 spinal cord segment.

An ophthalmic exam revealed very limited vision with bilateral diffuse retinal thinning, focal corneal dystrophy, and a focal incipient cataract in each eye. The corneal dystrophy and cataracts were considered likely to be congenital/hereditary. The retinal thinning was attributed to chronic anemia, though congenital/inherited progressive retinal atrophy could not be completely ruled out.

The dog was treated with a blood transfusion and prednisone (2 mg/kg/day). However, subsequent evaluations showed no improvement in neurologic signs, a progression of anemia, and increased respiratory effort. Two weeks after the initial presentation, due to a poor prognosis and deteriorating quality of life, euthanasia was elected, and the dog was submitted for postmortem examination.

### Pathological examination

On gross postmortem examination, the oral mucosa, ocular conjunctiva, and subcutaneous tissue were diffusely pale pink to white. The dorsal aspect of the cervical spinal cord appeared flattened. No other significant gross abnormalities were observed.

Microscopically, the main lesions were observed in the spinocerebellar tracts, with substantial lesions in the cerebellum. Approximately 85% of the cerebellum was atrophied with marked thinning of the molecular, Purkinje cell, and granule cell layers (Fig 1A-D). Luxol fast blue staining highlighted the loss of myelinated nerve fibers in the cerebellar white matter (Fig 1B). Within the affected white matter were increased numbers of reactive astrocytes including gemistocytes and type I Alzheimer cells immunolabelled with glial fibrillary acidic protein (GFAP) (Fig 1E and 1F). Frequent blood vessels in the white matter had low numbers of perivascular macrophages containing pale gray cytoplasmic pigment interpreted as lipofuscin. In the brainstem, the olivary nucleus had multifocal vacuolation and gliosis with gemistocytes and increased microglial cells (Fig 2A-2D). The cerebrum had bilaterally symmetrical similar white matter loss in all examined lobes (temporal, parietal, frontal lobes), involving the corpus callosum, centrum semiovale, and deep portions of corona radiata (Fig 3A). Sporadic blood vessels in these areas had mild perivascular cuffs of lymphocytes, plasma cells, and histocytes. The superficial subcortical white matter and the internal capsule were relatively spared. The neuronal cell bodies in the cerebral cortex, hippocampus, caudate nucleus, and midbrain were spared. The mesencephalic aqueduct was moderately dilated.

**Fig 1.**
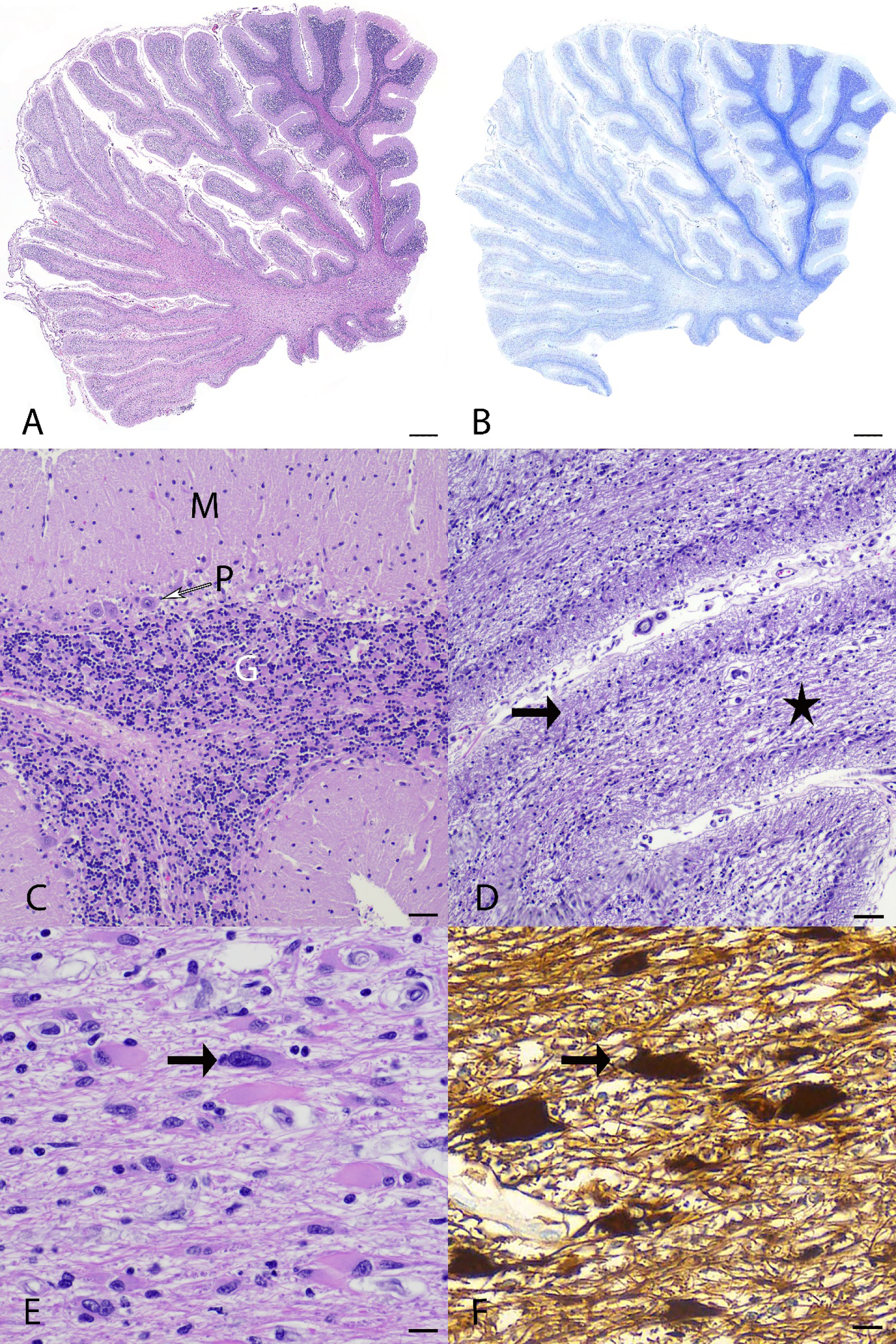
Histopathology of the cerebellum from a mixed-breed dog with spinocerebellar ataxia. **(A)** In the left side of figure, there is extensive full-thickness cortical degeneration with thinning of cerebellar folia. H&E stain. bar = 1000 µm. **(B)** Luxol fast blue stain highlights the loss of myelinated nerve fibers, characterized by loss of blue staining in the white matter, in those areas affected by cortical degeneration. Luxol fast blue stain. bar = 1000 µm. **(C)** In less severely affected areas of the cerebellum, all three cortical layers are preserved: molecular layer (M), Purkinje cell layer (P with an empty arrow), and granular layer (G). H&E stain. bar = 100 µm. **(D)** In severely affected areas of the cerebellum, the cortex has full-thickness degeneration with complete loss of Purkinje cells and marked thinning of the granule and molecular layers (arrow), while the white matter is thin, rarefied, and vacuolated (star). H&E. bar = 100 µm. **(E)** Cerebellar white matter contains plump pleomorphic astrocytes (arrow). H&E. bar = 50 µm. **(F)** Cerebellar white matter. Gemistocytes and Alzheimer type 1 astrocytes strongly immunolabeled for glial fibrillary acidic protein (GFAP) (arrow). Immunohistochemistry, bar = 50 µm.

**Fig 2.**
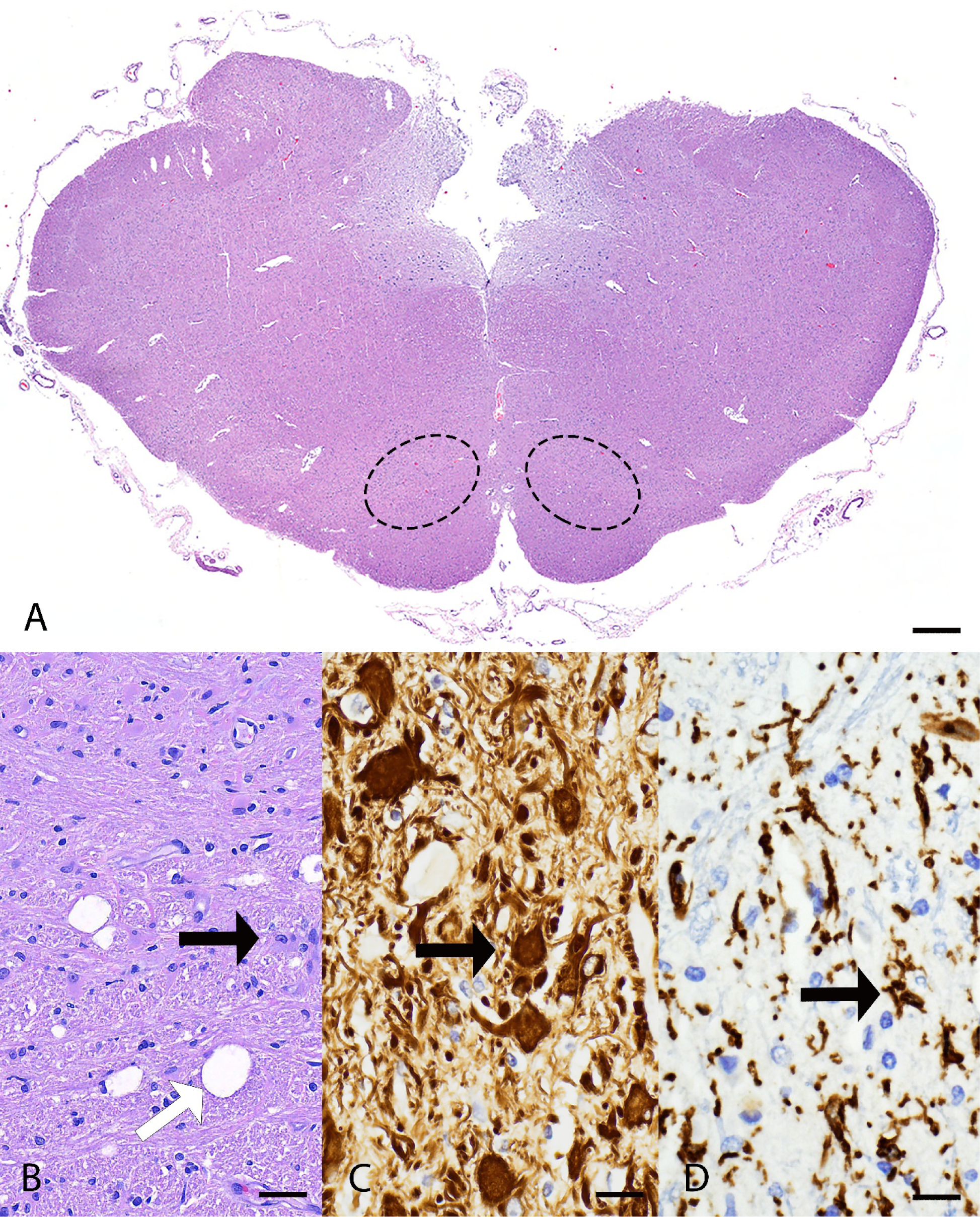
Histopathology of the brainstem from a mixed-breed dog with spinocerebellar ataxia. **(A)** In the brainstem, the olivary nuclei (highlighted with dashed ellipse) has multifocal vacuolation and gliosis. H&E stain. bar = 1000 µm. B-D higher magnification of olivaris nucleus. **(B)** There is vacuolation (white arrow) and moderate numbers of gemistocytes (black arrow) and increased microglial cells. H&E. bar = 50 µm. **(C)** Gemistocytes strongly immunolabeled for glial fibrillary acidic protein (GFAP) (arrow). Immunohistochemistry. bar = 50 µm. **(D)** There are increased numbers of activated microglial cells labeled with Ionized calcium-binding adaptor molecule 1 (IBA-1) (arrow). Immunohistochemistry. bar = 50 µm.

**Fig 3.**
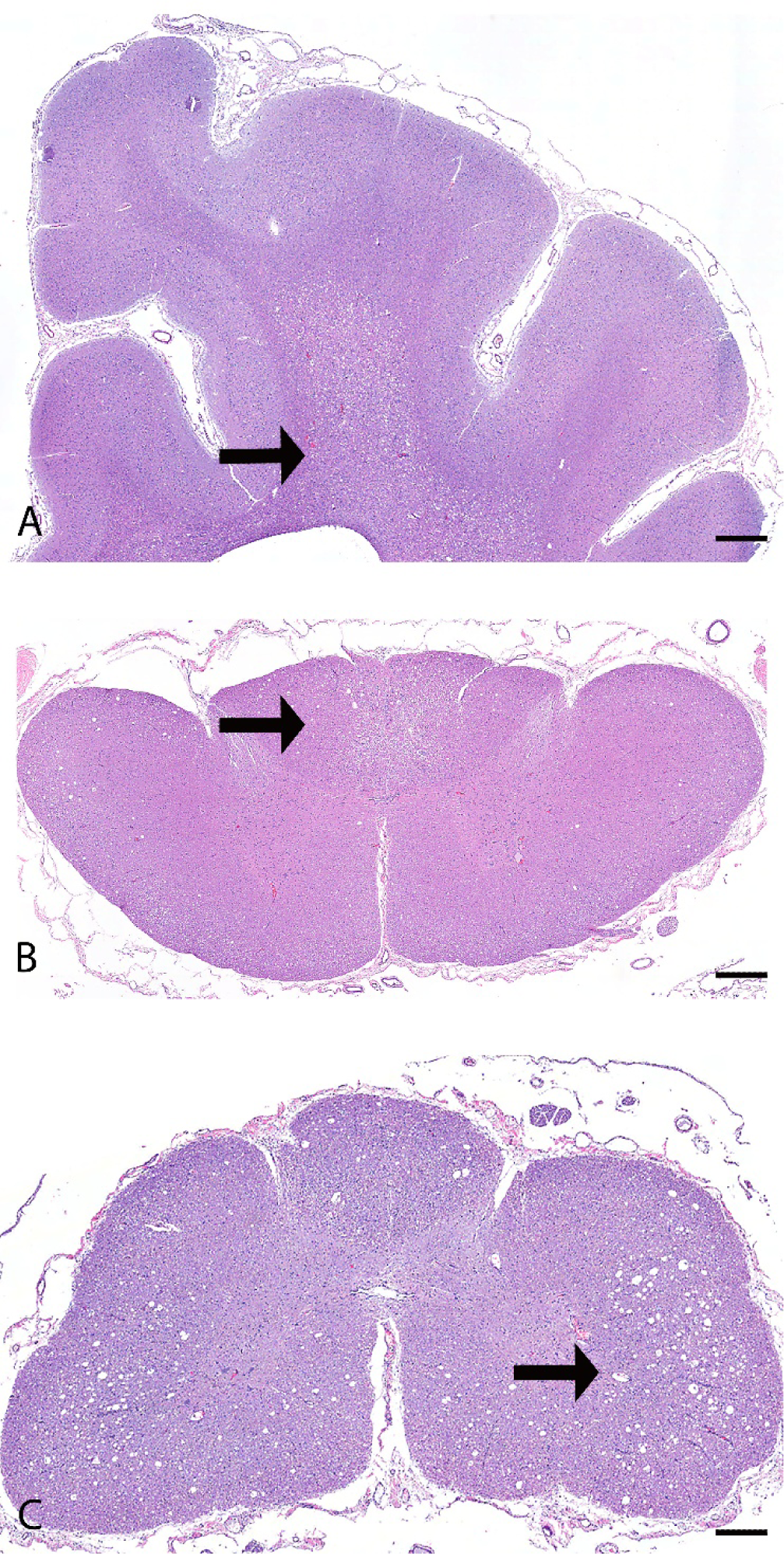
Histopathology of the cerebrum and spinal cord from a mixed-breed dog with spinocerebellar ataxia. **(A)** Cerebrum. The deep portion of the white matter is extensively vacuolated (arrow). H&E stain. bar = 1000 µm. **(B)** Cervical spinal cord. The white matter in the ascending pathway in the dorsal funiculus (arrow) and lateral funiculi is multifocally vacuolated. H&E. bar = 1000 µm. **(C)** Thoracic spinal cord. The white matter of the dorsal, lateral (arrow), and ventral funiculi is markedly vacuolated. H&E. bar = 1000 µm.

In the cervical spinal cord, the white matter in the ascending pathway, including the cuneate fasciculus and gracile fasciculus in the dorsal funiculus, and the dorsal and ventral spinocerebellar tracts in the lateral funiculi were multifocally vacuolated with digestion chambers, rare spheroids, and multifocal areas of gliosis (Fig 3B). The gracile fasciculus was particularly severely affected by loss of nerve fibers and scattered reactive astrocytes. The white matter in the descending pathway, mainly in the olivospinal and vestibulospinal tract in the ventral funiculi, was multifocally mildly vacuolated with digestion chambers. The neuronal cell bodies in the gray matter were spared. In the remaining spinal cord segments (thoracic and lumbar), the white matter of the dorsal, lateral, and ventral funiculi was markedly vacuolated, and frequently disrupted by gliosis, formation of digestion chambers and spheroids, and occasional reactive astrocytes (Fig 3C).

Both eyes had similar pathologic findings. The retina had multifocal areas of loss of ganglion cells and/or photoreceptor layer, and some areas had full-thickness atrophy with lack of distinction between all layers (Fig 4A). The retina also had multiple rosette-like structures, with separation in the photoreceptor layer or outer nuclear layer, interpreted as retinal folding secondary to degeneration (Fig 4A). The retina was segmentally detached with hypertrophy of the retinal pigmented epithelium (RPE) cells. The optic nerve had diffusely lost nerve fibers and was replaced by spindloid glial cells (interpreted as astrocytes) and thick dense collagenous septa (Fig 4B). The remaining structures including the cornea, uvea, and lens were unremarkable.

**Fig 4.**
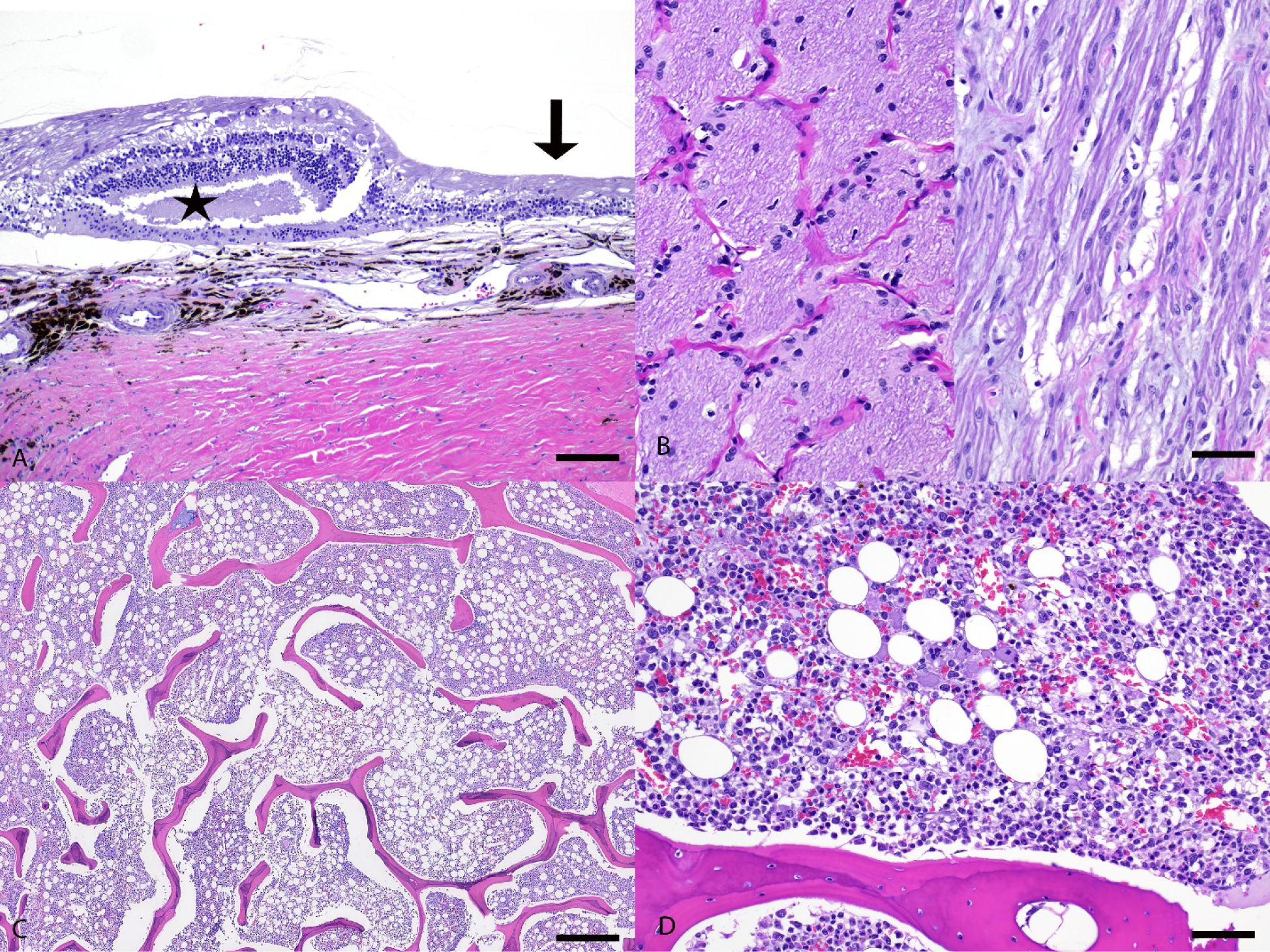
Histopathology of the eye and bone marrow from a mixed-breed dog with spinocerebellar ataxia. **(A)** The retina is disrupted by areas with rosette-like structures (star) and full-thickness atrophy (arrow) with lack of distinction between all layers. H&E stain. bar = 100 µm. **(B)** Left half demonstrates a normal canine optic nerve for comparison. Right half demonstrates the case dog’s optic nerve with loss of nerve fibers and replacement by glial cells. H&E. bar = 50 µm. **(C)** Bone marrow with approximately 60% cellularity (reference interval=25%-75%). H&E. bar = 500 µm. **(D)** Bone marrow with decreased myeloid to erythroid (M:E) ratio with the majority of cells being early erythroid precursors. H&E. bar = 100 µm.

In the bone marrow, the cellularity was approximately 60% (reference interval=25%-75%), and the myeloid to erythroid (M:E) ratio was decreased with the majority of cells being early erythroid precursors (Fig 4C and 4D). There was marked erythroid hyperplasia with left shift. These findings were not definitive for, but still supportive of, PIMA. There was marked granulocytic hypoplasia with marked left shift and toxic change, which was attributed to the bronchopneumonia mentioned below. There was also mild hemosiderosis and infiltration of a low number of plasma cells. The megakaryocyte lineage was unremarkable.

Additional co-morbid microscopic findings included neutrophilic bronchopneumonia with pleuritis and intralesional mixed bacterial colonies, and myocardial replacement fibrosis with sporadic individual cardiomyocyte degeneration and necrosis restricted to the papillary muscle of the left ventricle.

### Whole-genome sequence, variant filtering, and confirmation

None of the previously published canine spinocerebellar ataxia variants were observed in the case dog upon manual inspection using IGV (16). Therefore, the whole genome sequence of the case was first compared to the canine reference genome (UU_Cfam_GDS1.0_ROSY assembly), followed by comparison against 748 control genomes to identify variants present only in the case dog (Table 1, S2 Table). Variant prioritization focused on private, protein-altering variants present exclusively in either a heterozygous or homozygous state in the affected dog. VarElect was used with the search terms “spinocerebellar ataxia,” “anemia,” “paraparesis,” and “retinal thinning.” Genes with VarElect scores ≥ 10 were considered of particular interest.

**Table 1.** Results of variant filtering in a spinocerebellar ataxia-affected dog against 748 control genomes. VarElect search terms used were “spinocerebellar ataxia”, “anemia”, “paraparesis”, “retinal thinning;” only those scoring ≥ 10 were considered further.

| Filtering Step | Homozygous Variants | Heterozygous Variants |
| --- | --- | --- |
| All variants compared to reference | 2,586,116 | 3,213,087 |
| Private variants compared to 748 control canine genomes | 3,864 | 26,911 |
| Predicted protein-changing private variants | 17 | 175 |
| Located in functional candidate genes for similar phenotypes in other species, with VarElect score $\geq 10$ | 1 | 2 |

Variant filtering for a possibly dominant-acting mutation retained two heterozygous variants in functional candidate genes according to VarElect, both missense variants, one in *ERCC4* and one in *VWF*. Pathogenicity prediction tools PredictSNP (17) and MutPred2 (18), and others, classified the *ERCC4* variant (XP_038396243.1:p.Val81Ile) as likely benign, and it was therefore excluded (S3 Table). The *VWF* variant (NP_001002932.1:p.Arg1399His) yielded mixed predictions. It was classified as neutral approximately half of the time and pathogenic the other half; the latter all had lower confidence scores (S3 Table). In addition, there are no known associations between *VWF* and relevant neurological phenotypes.

While *VWF* variants are well-known to result in clotting disorders, this gene is only circuitously related to anemia (i.e., prolonged/heavy bleeding from the thrombopathia can result in an iron-deficiency anemia (19)); since “anemia” was a search term, this is likely driving the VarElect score for *VWF*. Therefore, both heterozygous variants were ruled out from further consideration.

The analysis identified only one homozygous protein-altering variant of “high” effect in a functional candidate gene. This variant was a homozygous 4-bp coding deletion (in exon 13 of 14) in *CLPX* (*caseinolytic mitochondrial matrix peptidase chaperone subunit*) and can be designated as XM_038580726.1:c.1723_1726del. This variant’s presence was confirmed in the case dog by visual examination in IGV; the four bp deletion was absent from a control dog (a Miniature American Shepherd, WGS generated from a different project, Fig 5). The *CLPX*:c.1723_1726del deletion is predicted to lead to a frameshift, inclusion of seventeen erroneous amino acids, and the creation of a premature stop codon (Fig 6), with predicted truncation of approximately 6.64% of the wild-type open reading frame (XP_038436654.1:p.Pro575CysfsTer18). The 4bp deletion variant is also absent from the most recently published Dog10K dataset comprising 1,987 genetically diverse canids (20).

**Fig 5.**
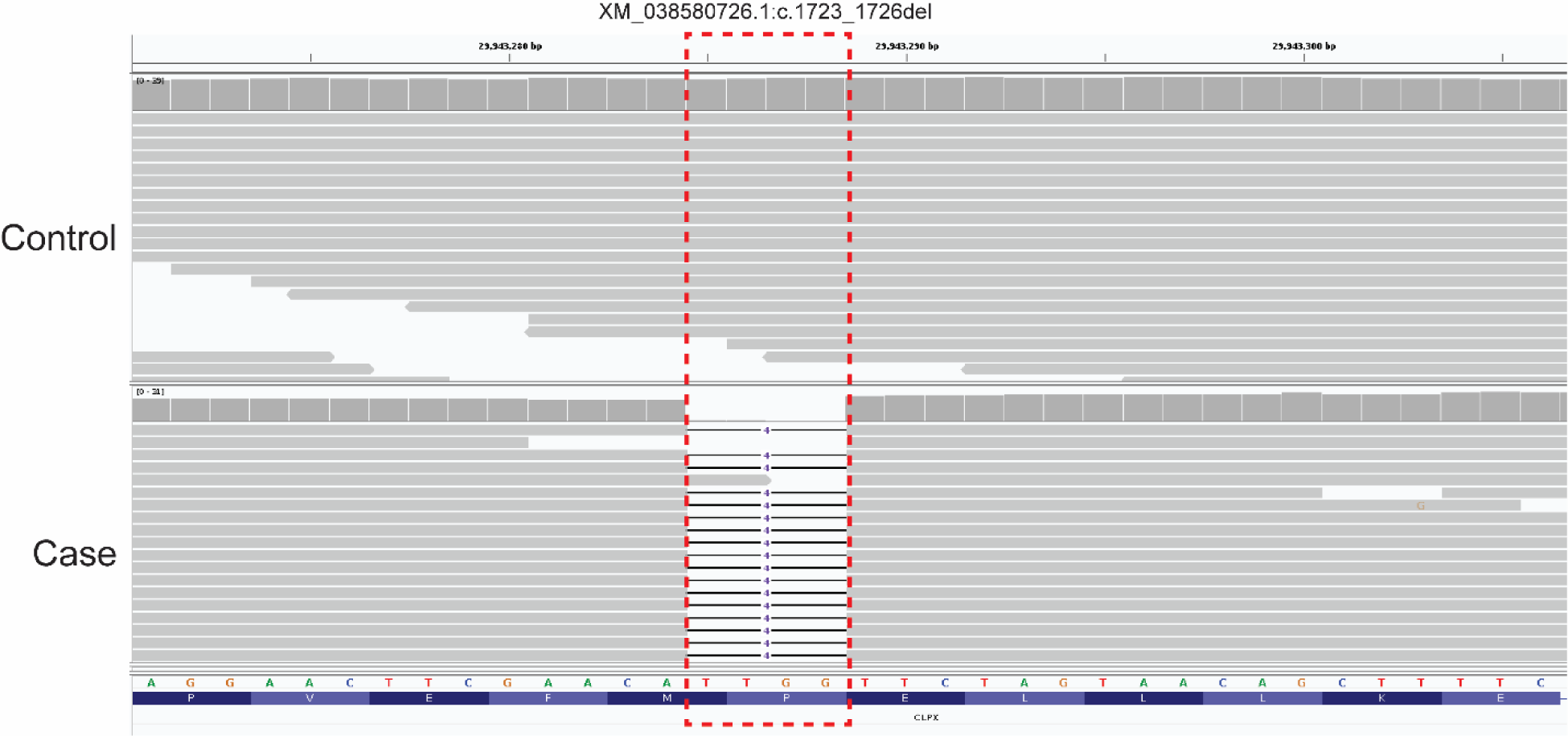
A 4bp deletion in canine *CLPX*. IGV screen shot of the short-read alignments from the affected dog (Case) and one unaffected dog (Control, a Miniature American Shepherd) demonstrating the 4bp deletion (CCAA; red box) in exon 13 (XM_038580726.1:c.1723_1726del). The sequence shown is in the reverse direction of the gene.

**Fig 6.**
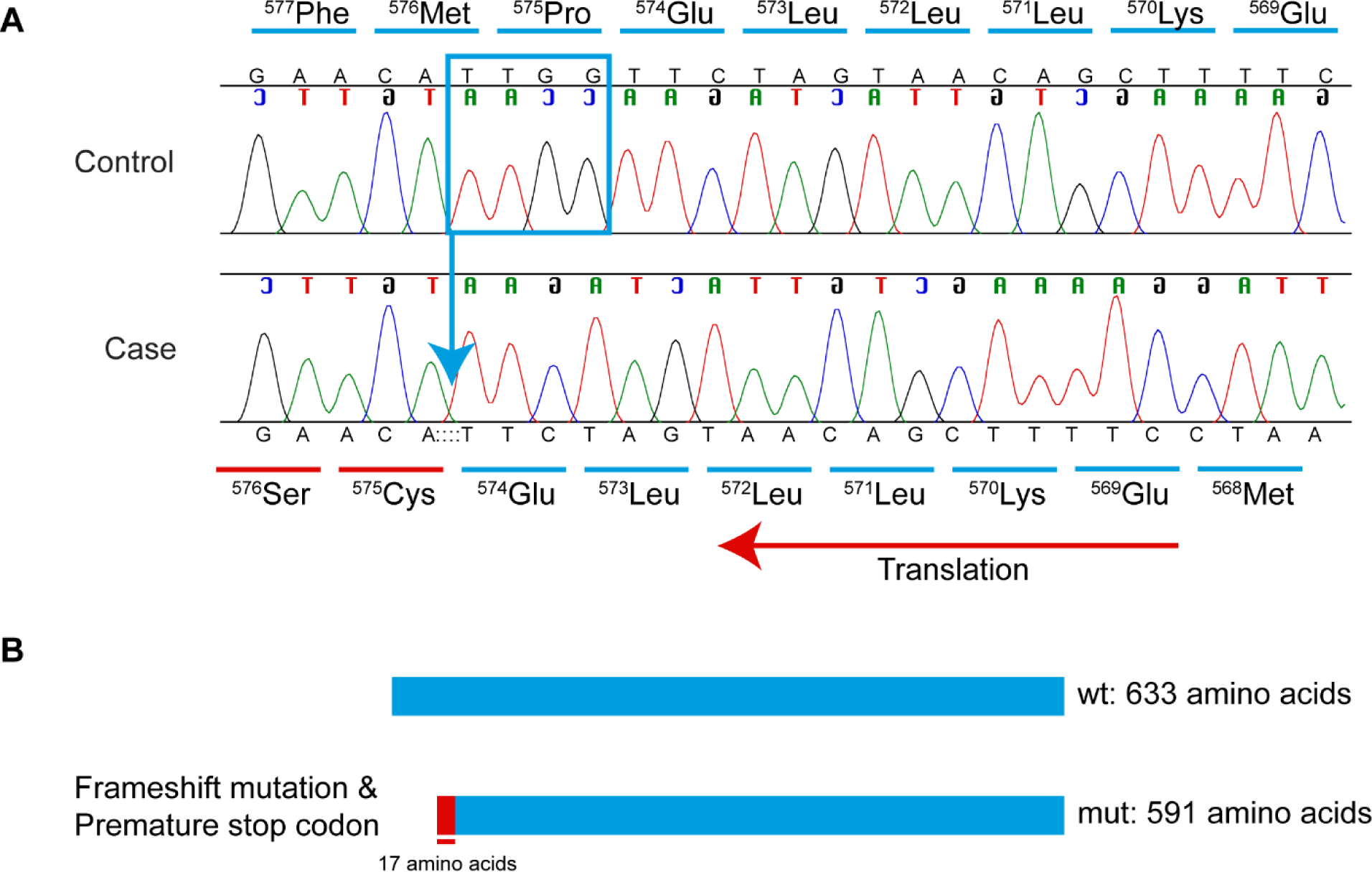
Sanger sequencing electropherograms demonstrating a 4bp *CLPX* deletion. **(A)** Chromatograms showing the *CLPX*:c.1723_1726del in the affected dog and a wild-type unaffected control Miniature American Shepherd. The correctly translated amino acid sequence is marked in blue above the control, and the frame-shifted codons are marked in red below the case. Complementary nucleotides representing the coding sequence are shown in reverse. **(B)** Schematic of wild-type and 4bp deletion variant-containing protein. The 4bp deletion is predicted to lead to a frameshift and a premature stop codon within 17 amino acids, likely truncating the resulting protein. The schematic is shown in the reverse direction, as the gene is on the reverse strand.

### Breed determination and principal components analysis

DNA from the case dog, reported as a Collie mix, was subjected to the commercially available Wisdom Panel breed determination test, as many SCA variants are known to be breed specific. This revealed the case mixed breed dog was comprised of approximately 30% Siberian Husky, 19% Rottweiler, 16% German Shepherd Dog, 8% American Staffordshire Terrier, and 8% Belgian Malinois, with much smaller contributions (all ∼6% or less) from American Pit Bull Terrier, Doberman Pinscher, Bulldog, Alaskan Malamute, Labrador Retriever, Weimaraner, and Soft Coated Wheaten Terrier. No Collie was detected in this dog. The WGS of the case was downsampled, using SNPs that overlap with the Axiom Canine HD 710K array. Principal component analysis (PCA) was then performed first with the five primary contributing breeds (Fig 7), and then again with all twelve contributing breeds (S4 Fig). To further investigate breed ancestry, PCA analysis was conducted again with the addition of Collies, Australian Shepherds, and Shetland Sheepdogs (S5 Fig), three closely related herding breeds, based on the report that the affected dog was a Collie mix. The case dog did not cluster directly with any of these three related herding breeds.

**Fig 7.**
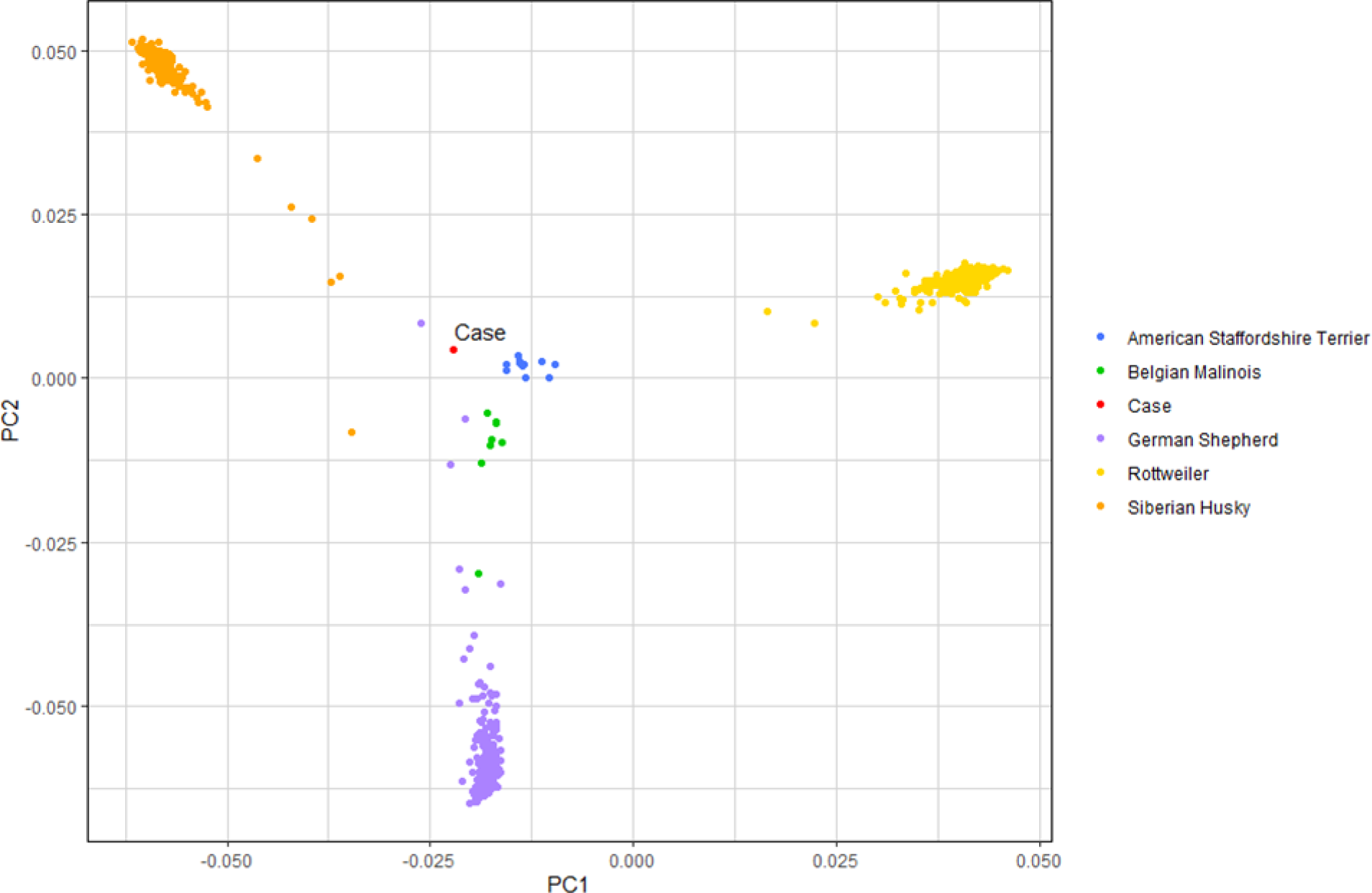
Principal component analysis (PCA). PCA of the genetic make-up of the case and five closely related dog breeds: American Staffordshire Terrier (n = 11), Belgian Malinois (n = 8), Case (n = 1), German Shepherd (n = 190), Rottweiler (n = 293), and Siberian Husky (n = 140).

## Discussion

We describe a *CLPX* variant that is associated with an SCA phenotype and nonregenerative anemia and provide a detailed characterization of the pathology. The clinical history and detailed pathology, revealing severe neurodegeneration predominantly involving spinocerebellar tracts, prompted further genetic investigation, leading to the identification of a homozygous four bp *CLPX* deletion present in the affected case. While nothing is known about the parents of the affected dog, they would each be obligate carriers for this deletion. It seems likely that the sire and dam were already related to each other, perhaps originally sharing this deletion variant in an identical-by-descent manner. This four bp deletion was absent from 2,735 other canine WGS.

CLPX is a subunit of a molecular chaperone involved in facilitating degradation of misfolded mitochondrial proteins, as part of a protein quality control (QC) system within the mitochondria; while this specific QC system normally plays a secondary role, its function becomes prominent when the cell is under cellular stress (21, 22). CLPX forms a hexameric ring to act alone or interact with CLPP peptidase forming a CLPXP complex (22). CLPX recognizes unstructured peptide degron, unfolds them, and translocates them to the CLPP compartment in the CLPXP complex where CLPP cleaves them into small peptides fragments (23). The evolutionary conservation of the CLPXP proteolytic complex from bacteria to humans underscores its fundamental role in mitochondrial protein quality control (22). *ClpP/ClpX* conditional knock out in mice caused the mitochondrial cristae structure to be disrupted in male germ cells. Moreover, *ClpP/ClpX* conditional knock out mice showed a decrease in mitochondrial membrane potential and inability to restore elevated reactive oxygen species (ROS) levels following oxidative stress in spermatocytes (24). CLPX also regulates heme synthesis – variants in this gene or related genes have been associated with erythropoietic protoporphyria 2 and anemia in humans (22, 25–27).

The deletion variant identified in this case is within a D2-small domain, which is essential for forming a tight interface with the D2-large domain of a neighboring subunit, that is known to provide binding energy to stabilize the functional assembly (28). Alteration in this domain may be associated with decreased protein stability. This *CLPX* frameshift variant is likely causative of this dog’s clinical signs and pathological findings. While the clinical phenotype of erythropoietic disorders has been previously associated with *CLPX* variants (25, 27), to date, neurologic signs have not been described with only a *CLPX* variant. However, related proteins in the CLPX protein pathway cause phenotypes similar or equivalent to SCA. A *CLPP* variant has been associated with human Perrault Syndrome, which includes phenotypic features of truncal and cerebellar ataxia (29). Variants in *AFG3L2*, a subunit of m-AAA protease that plays a role in degrading non-assembled and damaged protein in the mitochondria, is associated with SCA28 and Optic Atrophy 1 (30). Both the CLPP and ARF3L2 proteins are intimately involved in the same pathway with CLPX.

The diagnosis of SCA in this case was made based on the presence of progressive ataxia and neuropathology predominantly affecting the spinocerebellar tract. Histopathology revealed severe neurodegeneration in the cerebellum, spinal cord, deep cerebral white matter, and degeneration of the olivary nucleus, which plays a key role in coordinating signals between the cerebellum and spinal cord. The absence of primary inflammatory or infectious disease processes further supported a diagnosis of SCA. In addition to spinocerebellar tract degeneration, this case had optic nerve and retinal abnormalities, suggesting a broader, multisystem neurodegenerative process. According to the most recent four-group classification of canine hereditary ataxias proposed by Stee *et al*. (2023)(1), and given the involvement of the optic system and cerebral white matter in the present case, together with the dog’s widespread neurodegeneration of the spinocerebellar tracts, this case best fits the category multifocal degeneration with a predominant spinocerebellar component.

While neuropathology does not directly explain disease pathogenesis, patterns observed in hereditary ataxias with known genetic variants suggest that gene function often aligns with specific neuropathologies. In particular, there have been cases of canine SCA in which the causative gene is involved in mitochondrial function, and all of these have been classified in the category of multifocal degeneration with a predominant spinocerebellar component (1, 31–33). This supports the idea that mitochondrial dysfunction contributes to more widespread neurodegeneration, which is consistent with the *CLPX* deletion variant identified in this case. The significance of mitochondrial dysfunction in SCA is well-documented in humans, as mitochondrial function impairment leads to oxidative stress and energy deficits, particularly in high-energy-demanding tissues, especially the central nervous system (CNS) (34). In humans, mitochondrial ataxia is more commonly a multisystem disorder, rather than affecting a single tissue; the commonly affected tissues include CNS, retina, heart, endocrine glands, and bone marrow (34).

Initially, the retinal thinning observed on ophthalmic examination was attributed to anemic retinopathy. However, given the concurrent optic nerve degeneration, multiple retinal folds/rosette-like structures, and identification of a *CLPX* gene deletion variant, it is more plausible that the ophthalmic changes are part of a broader neurodegenerative process associated with the *CLPX* variant. Ophthalmic involvement has been reported in various subtypes of SCA in humans. For example, SCA7, an autosomal dominant SCA, is characterized by progressive retinal degeneration and vision loss in addition to cerebellar ataxia, which is histologically characterized by loss of the photoreceptor and ganglion cell layers and thinning of all retinal layers (35, 36). Several types of autosomal recessive cerebellar ataxia also have ocular manifestations including retinitis pigmentosa, chorioretinopathy, and optic neuropathy (37). These conditions are associated with variants in genes involved in diverse biological processes including vitamin E metabolism (gene: *TTPA*), breakdown of endocannabinoid neurotransmitters in the CNS (gene: *ABHD12*), chloride channels (gene: *CLCN2*), and mitochondrial poly (A) polymerase (gene: *MTPAP*) (37). Mitochondrial diseases have a tendency to affect metabolically active tissues, and ocular involvement is present in over half of all human patients with mitochondrial disease, variably affecting the retina, optic nerve, and extraocular muscles (38). Similarly, in dogs with hereditary ataxia, clinical signs suggestive of visual loss such as reduced or absence of menace response have been reported in many cases (1). However, other than a few cases of lysosomal storage diseases, histopathology of the eyes are not commonly reported in canine cases (1, 39, 40). In the present case, the retinal and optic nerve degeneration explains the dog’s poor vision and aligns with the patterns of tissue involvement seen in other mitochondrial disorders.

This canine patient also had severe non-regenerative anemia, which was initially suspected to be PIMA based on the clinical history of ineffective erythropoiesis and previous response to steroid treatment. Even though rubriphagocytosis, a common feature of PIMA, was not identified in bone marrow histology in this dog, the clinical and histopathological findings of chronic non-regenerative anemia, erythroid hyperplasia with left shift, and hemosiderosis are supportive of PIMA (41). However, the identification of the *CLPX* variant raises the possibility of mitochondrial dysfunction being responsible for the anemia. Mitochondrial dysfunction has been implicated in the development of anemia through several mechanisms (22, 42–44). *CLPX* plays an important role in cellular heme synthesis, and, significantly, *CLPX* variants have been previously associated with erythropoietic disorders (25). CLPX regulates aminolevulinic acid synthase 2 (ALAS2), the first enzyme in heme production, by activating it via its unfoldase activity and by degrading it via transporting it to CLPXP (27). CLPX is also required for iron utilization for heme synthesis during erythroid differentiation (22). Variants in the *ALAS2* gene, encoding ALAS2 (regulated by CLPX) can cause hereditary sideroblastic anemia in humans (42). A naturally occurring *CLPX* variant associated with an erythropoietic disorder has been reported in humans; a heterozygous missense variant in *CLPX*, which partially inactivates the protein by inhibiting ClpXP complex formation, has been associated with erythropoietic protoporphyria in a human family (45). This disruption leads to increased ALAS activity, because CLPX presumably retains its ability to activate ALAS while losing its ability to degrade it (27, 45). While to our knowledge a *CLPX* variant has not been associated with naturally occurring anemia, it can be speculated that the truncating *CLPX* variant in our case may have caused more severe loss of function of the protein compared to the human missense variant, resulting in loss of both the ALAS degrading and activating function, leading to anemia. This theory is supported by findings that *CLPX* knockout in murine erythroid cells reduced hemoglobin and heme synthesis, and that *CLPX* knockout in zebrafish similarly resulted in anemia (26, 45). Additionally, chronically increased oxidative stress caused by the *CLPX* variant (24) could have predisposed the dog to the development of immune-mediated anemia. Oxidative stress is known to contribute to the development or exacerbation of immune-mediated diseases by leading to macromolecule modification, generation of new auto-antigen, and inducing pro-inflammatory response (46, 47). Elevated ROS levels can lead to oxidative modifications of red blood cell (RBC) components, making them antigenic and triggering autoimmune responses (48). For instance, studies have shown that increased ROS levels in RBCs correlate with the production of autoantibodies against these cells in New Zealand black mice, which are prone to autoimmune hemolytic anemia (48). The myocardial fibrosis and focal cardiomyocyte necrosis seen in the case dog may reflect secondary hypoxic injury associated with the severe anemia, although a direct contribution of the *CLPX* variant to cardiac pathology cannot be excluded.

Western blot analysis (S6 Appendix) confirmed that CLPX expression was still present in the affected dog compared to control (S7-S9 Fig). CLPX is highly conserved between humans and dogs, with 98.42% sequence identity according to Protein Blast. The four-base pair *CLPX* deletion is only predicted to truncate the amino acid chain by approximately 6.64%; this together with the western blot results suggest that the transcript is not removed via nonsense-mediated decay. Since protein expression is comparable to the control, the deletion likely disrupts an important functional domain rather than affecting the overall protein stability. This supports the idea that appropriate levels of functional CLPX are necessary for normal mitochondrial activity. Homozygous knock-out *Clpx ^−/−^* mice die very early in embryonic development (49). Because the variant in our case dog was very close to CLPX’s C-terminus, truncating only a small portion of the protein, and as expression of CLPX was verified by our western blot findings, it is reasonable to conclude that CLPX must have retained enough function to be compatible with complete embryonic development.

Once WGS was generated for the case, it became evident that the dog was not primarily Collie, as initially presumed. This spurred commercial breed testing, which indicated that the case was a highly mixed breed dog, with no Collie ancestry. This finding highlights the limitation of visual breed identification in mixed breed dogs and underscores the importance of molecular breed determination for accurate classification. Previous work has clearly indicated that assigning breed via visual inspection only is fraught with inaccuracies (50, 51). Accurate breed information is particularly important when investigating genetic diseases, as many disease-causing variants are restricted to certain breeds; therefore, clinicians and pathologists must always view reported breed, especially in mixed-breed animals, with some level of skepticism, and when possible, confirm ancestry through genetic testing.

A major limitation of the present study is that it describes only a single affected animal; while there were reportedly affected littermates, none were available for further investigation. Likewise, DNA was not available from either parent to verify their heterozygosity for the 4bp deletion. It is possible that other modifying variants not detected in this pipeline could be contributing to this dog’s phenotypic spectrum; however, it is unlikely any other significant, single-gene variants would be causative, barring large structural rearrangements which are not easily detected using short-read WGS. Additionally, an antemortem bone marrow aspirate could have provided more detailed and reliable information about the nature of the anemia.

## Conclusion

This study presents a case of SCA categorized as multifocal degenerations with a predominant (spino)cerebellar component in a young mixed breed dog with retinal and optic nerve degeneration and nonregenerative anemia. A 4bp frameshift variant was detected in the gene *CLPX*. This is the first case report of SCA associated with a naturally occurring *CLPX* variant in any species. This case highlights how integration of clinical, pathological, and genetic findings can uncover novel disease mechanisms and candidate genes; furthermore, it demonstrates that spontaneous animal models can provide new insights which can then be translated back to people. While traditional neuropathologic classification remains useful, it may not fully reflect underlying pathogeneses. As more causative variants are discovered in dogs, incorporating genetic information into classification frameworks will be essential for advancing our understanding of SCAs in dogs and improving diagnostic precision.

## Materials and methods

### Clinical examination and ethical statement

A 1-year-and-11-month-old male neutered dog, reported as a Collie mix, was referred to the Louisiana State University Veterinary Teaching Hospital (LSU VTH) for weakness—particularly of the pelvic limbs— and severe non-regenerative anemia. The dog underwent clinical examination, including both neurologic and ophthalmic exams. Bloodwork performed included a complete blood count (CBC), serum chemistry analysis, saline agglutination test, and rapid vector-borne infectious disease panel. A genetic test for degenerative myelopathy (15) was also carried out, since littermates were reported to likewise be affected with pelvic limb weakness. Due to progressive clinical signs and failure to respond to treatment, humane euthanasia was elected under the care of the supervising veterinary clinician.

The work described in this manuscript involved a non-experimental diagnostic necropsy on an animal that was euthanized based on veterinary recommendation and owner consent. Therefore, institutional ethical approval was not required. The postmortem examination was conducted with the owner’s informed consent and followed standard veterinary necropsy procedures.

### Pathological examination

The dog was submitted for postmortem examination to the Louisiana Animal Disease Diagnostic Laboratory (LADDL), Louisiana State University (Baton Rouge, LA, USA). Samples from all major organs, including the liver, heart, spleen, lung, kidneys, brain, small and large intestines, stomach, bone marrow, urinary bladder, spinal cord, eyes, and adrenal glands, were fixed in 10% neutral-buffered formalin for 24 to 48 hours. Following fixation, tissues were processed routinely, embedded in paraffin, and 4μm sections of formalin-fixed paraffin-embedded (FFPE) tissues were cut and stained with hematoxylin and eosin (H&E) for histologic evaluation according to standard laboratory procedures. Selected sections of the brain and spinal cord were stained with Luxol fast blue and selected sections of the brain only were immunolabeled for glial fibrillary acidic protein (GFAP) and ionized calcium-binding adaptor molecule 1 (IBA-1). Briefly, 4μm sections of FFPE tissue sections were mounted on positively charged Superfrost® Plus slides (VWR, Radnor PA) and subjected to IHC using the automated BOND-MAX and the Polymer Refine Detection kit (Leica Biosystems, Buffalo Grove, IL). Following automated deparaffinization, heat-induced epitope retrieval (HIER) was performed using a ready-to-use citrate-based buffer (pH 6.0; Leica Biosystems) for anti-Iba-1 at 100 °C for 20 min before incubation with the primary antibody. For GFAP, no retrieval was performed. Sections were then incubated with the primary antibodies (anti-Iba-1 at 1:2,000 [Wako Chemicals, 019-197-41] and anti-GFAP [Dako, Z0334] at 1:500) for 30 min at room temperature, followed by a polymer-labeled goat anti-rabbit IgG coupled with horseradish peroxidase (Leica Biosystems) for 8 minutes at room temperature. 3,3’-diaminobenzidine tetrahydrochloride (DAB) was used as the chromogen (10 minutes), and counterstaining was performed with hematoxylin. Slides were mounted with a permanent mounting medium (Micromount®, Leica Biosystems).

### DNA Extraction

Fresh-frozen cardiac tissue (obtained at necropsy) was shipped from LSU to the Purdue Canine Genetics Laboratory. Because the tissue was obtained via necropsy, no IACUC approval was required. DNA was extracted from the cardiac tissue using the Qiagen DNeasy Blood and Tissue kit, following the manufacturer’s protocol.

### Whole-genome sequencing, alignment, and variant calling

All genomic locations and positions are in reference to the UU_Cfam_GSD_1.0_ROSY/CanFam4.0 genome assembly. DNA from the case dog was subjected to whole-genome sequencing (WGS) at Azenta Life Sciences (South Plainfield, NJ) using Illumina NovaSeq 150bp paired-end reads, with an average of 35.9x coverage. Quality control, mapping, and variant calling of reads was performed using the Whole Animal Genome Sequencing (WAGS) pipeline (52). Briefly, reads were aligned to the UU_cfam_GSD_1.0_ROSY reference genome using the Burrows-Wheeler Alignment Tool (53). Variant calling was then executed with Genome Analysis Toolkit (GATK) following Best Practices (54), and Ensembl’s Variant Effect Predictor (VEP) (55) was used to predict the consequence of variants.

### Variant filtering

As an initial screening approach, IGV was used to manually examine the case dog’s WGS for any published canine variants associated with spinocerebellar ataxia as curated by Online Mendelian Inheritance in Animals (OMIA) (accessed July 2025) (56) or previously described (1).

Next, genetic variants from this case were compared to 748 control dog genomes from an existing internal WGS database of genetically diverse dog breeds. Variants private to the case were identified with BCFTools1.17 (57) under the assumption that no other dogs in the existing WGS database carried the same disease-causing allele; next, the private variants were prioritized based on predicted impact (“high” or “moderate”) on the resulting transcript with VEP together with the Dog10K annotation for UU_Cfam_GSD_1.0_ROSY genome reference assembly (58). The resulting gene name of each “high” or “moderate” variant was then filtered through VarElect (59) which gives a score to each gene as an indication of the strength of association between that gene and a list of phenotypic search terms, in this case, “spinocerebellar ataxia”, “anemia”, “paraparesis”, or “retinal thinning”. Genes with a VarElect score ≥ 10 were considered likely to cause disease and were further investigated.

### In-silico predictions

The impact of the frameshift deletion on the resulting transcript was predicted using the ExPASy Translate tool (60). The DNA sequence of canine *CLPX* was retrieved from NCBI and the sequence with the 4bp deletion (XM_038580726.1:c.1723_1726del) was translated to amino acids and compared to the wild-type protein sequence. The molecular weight of the predicted amino acid sequence containing the 4bp deletion was calculated with ExPASy Compute pI/Mw (60).

### Primer design and targeting genotyping

Primers were designed to amplify over the 4bp deletion for further confirmation via Sanger sequencing. Primer3Plus (61) (https://www.primer3plus.com/index.html) was used to design primers to amplify a 584bp (or 588bp in case of the wild-type allele) region surrounding the 4-bp *CLPX* deletion in the case dog and one control (a Miniature American Shepherd). PCR was used to amplify the 584bp region from genomic DNA using KOD Xtreme Hot Start DNA Polymerase (Millipore-Sigma, Burlington, MA) and primers F-5’-TTAGGGGGAAAAAGTGCAGGC-3’ and R-5’-GTGCATGGTGATCTAAGAAGGC-3’ using an annealing temperature of 60°C and standard conditions. Amplicons were treated with ExoSAP-IT PCR Cleanup Reagent (Thermo Scientific, Waltham, MA) and submitted for Sanger sequencing (Eurofins Genomics, Louisville, KY). Sanger sequences were analyzed using Sequencher5.4.6 (GeneCodes, Ann Arbor, MI, USA).

### Breed determination and principal component analysis

Because the case was reported as a Collie mix and had come through a rescue making this heritage dubious, DNA from this dog was subjected to the commercially available Wisdom Panel breed determination test in order to determine the breed composition of the case. Based on these results, the case’s WGS was downsampled by extracting 710K SNPs (equivalent to the Axiom CanineHD SNP array) and merged with an internal database of canine SNP data (from internal studies, collaborator data, and publicly available data) from more than 10,000 dogs. The following dogs were then extracted from this database: the case dog (n = 1), Alaskan Malamute (n = 9), American Bulldog (n = 2), American Pitbull Terrier (n = 6), American Staffordshire Terrier (n = 11), Australian Shepherd (n = 294), Belgian Malinois (n = 8), Collie (n = 5), Doberman Pinscher (n = 77), French Bulldog (n = 101), German Shepherd (n = 190), Labrador Retriever (n = 1,434), Rottweiler (n = 293), Shetland Sheepdog (n = 63), Siberian Husky (n = 140), and Weimaraner (n = 2). With this set of SNP data, basic quality control filtering was performed in PLINK 1.9 (62). Dogs were eliminated if they were missing more than 10% of SNP calls. SNPs were eliminated due to low call rate (< 90%), low minor allele frequency (< 0.05), and extreme deviation from Hardy Weinberg equilibrium (p-value < 0.000001); 2,636 dogs and 62,209 SNPs remained for analysis. PCA analysis was carried out using Genome-wide Complex Trait Analysis (GCTA) (63) software. The Collie, Australian Shepherd, and Shetland Sheepdog were included in follow-up analysis because the case dog was initially reported to be a Collie mix, and the Australian Shepherd and Shetland Sheepdog are the breeds most closely related to Collies (64) for comparison.

## Acknowledgements

The authors thank Drs. Jocelyn Lee, Kylie Landry, and Patty Lathan at LSU VetMed for their role in the clinical management of this case and for sharing case information that supported the preparation of this report. We thank the additional pathologists consulted in this case who made significant contributions to the diagnosis: Dr. Porter (Texas A&M), Dr. Rissi (UGA), Dr. Koehler (Auburn), and Dr. Langohr (SANOFI). We thank Dr. Shannon Dehghanpir for her expertise in bone marrow evaluation, and the personnel from the Histology and Immunohistochemistry section at the Louisiana Animal Disease Diagnostic Laboratory. The authors are also grateful to Wisdom Panel, Science and Diagnostics, a division of Mars Petcare, for donating their breed determination test. We acknowledge USDA NAHLN and the Kenneth Burns Endowed Chair in Veterinary Medicine support at LSU Vet Med for funding digital slide scanners.

## Data Availability

Raw whole-genome sequencing data for the affected dog is available from SRA under the accession number PRJNA1290392 and sample accession number SAMN49918932.

## Author Contributions

Conceptualization: BSD, JMB, MCC, KJE, JL

Data curation: BSD, JMB, NJK, AJ, KJE, JL

Formal analysis: BSD, JMB, NJK, MCC, KJE, JL

Funding acquisition: KJE

Methodology: BSD, JMB, ADM, KJE, JL

Resources: BSD, KJE, JL

Supervision: KJE, JL

Visualization: BSD, JMB, JL

Writing – original draft: BSD, JMB, NJK, KJE, JL

Writing – review & editing: BSD, JMB, NJK, MCC, AJ, ADM, KJE, JL

## Supporting information

**S1 Table. Complete blood count results from the case dog.**

**S2 Table. Variant analysis.** Excel workbook (multiple sheets) detailing genetic variants identified and prioritized in case dog.

**S3 Table. Protein pathogenicity predictive software results for two heterozygous variants.** List of heterozygous variants with VarElect scores ≥10 and predicted pathogenicity scores from multiple programs.

**S4 Fig. Principal component analysis (PCA) of the genetic make-up of the case and twelve different dog breeds.**

**S5 Fig. Principal component analysis (PCA) of the genetic make-up of the case and fifteen different dog breeds.**

**S6 Appendix. Western blot analysis methods.**

**S7 Fig. Representative western blot results of CLPX in cardiac tissues from an unaffected control dog and the case dog.**

**S8 Fig. Original untrimmed image of the Western blot analysis performed to detect CLPX in the cardiac muscle tissue from the case dog and an unaffected control dog.**

**S9 Fig. Original untrimmed image of the Western blot analysis performed to detect β-actin in the cardiac muscle tissue from the case dog and an unaffected control dog.**

